# Mechanical Properties of the Developing Brain in a Model of Fetal Alcohol Spectrum Disorders and Relationships to Perineuronal Net Integrity

**DOI:** 10.1101/2025.10.07.680991

**Authors:** L. Tyler Williams, Katrina A. Milbocker, Eva Smith, Cassandra P. Bodner, Diego A. Caban-Rivera, Matthew D. J. McGarry, Elijah E. W. Van Houten, Anna Y. Klintsova, Curtis L. Johnson

**Author notes:** Correspondence: Curtis L. Johnson, PhD 540 S. College Avenue, Suite 109C Department of Biomedical Engineering University of Delaware Newark, DE 19713. authors contributed equally (L. Tyler Williams, Katrina A. Milbocker).

## Abstract

Neuroimaging is a useful tool for examining altered neurodevelopmental trajectories in fetal alcohol spectrum disorders (FASD). FASD affects 1 in 20 infants in the United States with higher prevalence in specific regions across the globe. Advanced neuroimaging methods, such as volumetric morphometry and diffusion-weighted imaging, are critical for determining the effectiveness of interventions that support neurodevelopment in FASD. In this study, we introduce the use of magnetic resonance elastography (MRE), a cutting-edge neuroimaging technique used to measure the mechanical properties of brain tissue, to assess the impact of alcohol exposure and combined exercise and environmental complexity intervention on neurodevelopment in a rat model of FASD. Our results indicate that brain stiffness is reduced in juvenile alcohol-exposed rats which is recovered to baseline by adulthood, and damping ratio increases in all rats with age. Additionally, we quantified cortical perineuronal net (PNN) density which follows similar trends to shear stiffness and damping ratio, suggesting MRE may be an effective method for noninvasively monitoring FASD progression related to extracellular matrix integrity.

## 1. Introduction

Fetal alcohol spectrum disorders (FASD) encompass a group of neurological conditions resulting from prenatal alcohol exposure. Acute alcohol insult delays the trajectory of neurodevelopment, leading to persistent cognitive deficits across adolescence (May et al., 2015). Neurodevelopmental delay due to FASD has been monitored using conventional longitudinal neuroimaging techniques, including volumetric morphology, diffusion-weighted imaging, and functional near-infrared spectroscopy, which have been used to uncover persistent changes to brain parenchymal growth, cortical organization and thinning, myelination, and maturation of vasculature in the brains of children and adolescents (Norman et al., 2009; Riley et al., 1995; Rockhold et al., 2023; Smiley et al., 2015; Sowell et al., 2001; Zhou et al., 2011). While MRI has been used to identify FASD-specific anomalies in brain structure and function, short-term microstructural changes that precede and drive macroscale structural damage are more challenging to measure, impeding intervention development and disease progression monitoring (Hoyme et al., 2016; Popova et al., 2018).

Magnetic resonance elastography (MRE) is a novel imaging technique that uses applied shear waves to estimate mechanical properties of tissues, akin to palpating tissue noninvasively (Manduca et al., 2001; Muthupillai et al., 1995). Mechanical properties measured through MRE include shear stiffness and damping ratio which can reflect microstructural composition and organization (Manduca et al., 2021; Sack, 2023; Sack et al., 2013). MRE is the gold-standard method for staging liver fibrosis in alcohol-use disorders and other related diseases (Kennedy et al., 2018). When applied to the brain, it has shown promise in evaluating tissue integrity as it is sensitive to small variations in tissue microstructure which may precede volumetric changes due to neuropathology (Hiscox et al., 2016; Murphy et al., 2019). In the adult brain, lower shear stiffness has been observed in several neurodegenerative diseases including Alzheimer’s disease, multiple sclerosis, and Parkinson’s disease (Hiscox, Johnson, McGarry, Marshall, et al., 2020; Lipp et al., 2013; Sack, 2023; Sandroff et al., 2017). Furthermore, shear stiffness decreases with natural aging, reflecting a steady decline in brain health into old age (Hiscox et al., 2021). Damping ratio, or other measures describing relative viscosity, is reported less frequently than shear stiffness, yet it may be more sensitive to individual differences and has been associated with certain cognitive performance measures, making it a valuable parameter in assessing overall tissue health (Delgorio et al., 2023; Hiscox, Johnson, McGarry, Schwarb, et al., 2020; Sack, 2023; Schwarb et al., 2017; Williams et al., 2024).

While MRE has most commonly been applied to the adult and older adult brain, there are also many applications in the developing brain that warrant further investigation. It is well established that the developing brain undergoes numerous and complex microstructural changes (Juraska & Drzewiecki, 2020; Lebel et al., 2008; Makropoulos et al., 2016). Like the aging brain, these microstructural changes have macroscale effects on tissue mechanical properties and may be disrupted by pathology and damage (Johnson & Telzer, 2018). However, developmental MRE studies are limited, and results vary on mechanical property differences between children and adults (Hiscox et al., 2018; McIlvain et al., 2018, 2022). There are even fewer studies on development with neurodegenerative disease, but one found lower stiffness in children with cerebral palsy (Chaze et al., 2019).

Similar trends have been found throughout the preclinical MRE literature. Brain stiffness decreases in rodent models of Alzheimer’s disease, multiple sclerosis, and stroke (Bertalan et al., 2019; Freimann et al., 2013; Munder et al., 2018; Murphy et al., 2012; R. V. Silva et al., 2021; Wang et al., 2020). Preclinical studies on rodent development show increased brain stiffness with increased myelination and cytoskeletal linking (Guo et al., 2019; Pong et al., 2016). Our previous pilot study using a rat model of FASD captured a transient reduction to forebrain shear stiffness and damping ratio in alcohol-exposed rats during late adolescence, on postnatal day (PD) 42, that was resolved by young adulthood on PD 70. We suspect that this may be due to the characteristic delay in myelination that occurs in this model and clinically. However, this correlation was not observed, suggesting that other microstructural factors contribute to longitudinal changes in brain stiffness in this model. Moreover, exercise had a transient effect on mechanical properties that may have corrected for delayed neurodevelopmental trajectories (Milbocker et al., 2024). These findings suggest the use of MRE in preclinical models of FASD can reveal sensitivity to disrupted microstructure and intervention. To understand this relationship, further study is needed during the early stages of development combined with correlations to additional microstructural components through histology.

Alcohol is a known teratogen – a process that can cause birth defects or abnormalities during fetal development – that disrupts neuronal survival and differentiation, neurite outgrowth, glial organization, cell adhesion, and circuit connectivity during critical periods which contribute to delayed or disorganized neurodevelopment (Dannenhoffer et al., 2022; Lasek, 2016; Licheri & Brigman, 2021; Milbocker et al., 2022; Miñana et al., 2000). Several factors including circuit connectivity and oligogenesis have already been linked to mechanical property variations in preclinical studies (L. Liu et al., 2023; Milbocker et al., 2022, 2024). Perineuronal net (PNN) integrity is another factor that may prove valuable in assessing FASD progression and may be assessed *in vivo* with MRE. PNNs are a major component of the brain extracellular matrix that primarily surround inhibitory parvalbumin interneurons and contribute to local brain tissue mechanical properties (Bergs et al., 2024). Their integrity is susceptible to prenatal alcohol exposure and damage persists long term, indicating that cytoskeletal components are most likely affected by prenatal alcohol (Lewin et al., 2018; Saito et al., 2019). Moreover, several reports indicate that the organization and maturation of *cortical* parvalbumin interneurons is significantly affected by prenatal alcohol exposure, likely related to maladaptive PNN formation and ultimately contributing to lack of inhibitory control (Licheri et al., 2023; Marguet et al., 2020; Smiley et al., 2015). MRE is sensitive to extracellular matrix changes in mice and tissue stiffness has been has been reported to correlate with PNN integrity in a model of experimental autoimmune encephalomyelitis (Batzdorf et al., 2022; R. V. Silva et al., 2024). Therefore, given the mechanosensitivity and integral nature of PNNs to maintaining brain microstructure, MRE may provide a method of noninvasively monitoring PNNs in the FASD brain.

In this study, we utilize MRE to explore the effects of alcohol exposure, exercise, and environmental complexity on mechanical properties during the juvenile (PD 25-29) and young adult (PD 80-84) phases. From our previous work on adolescent rats (Milbocker et al, 2024), we expect alcohol-exposed, juvenile rats to exhibit softer brain tissue which is recovered to normal levels by adulthood. Exercise intervention was shown to have a positive effect on shear stiffness, allowing alcohol-exposed rats to “catch up” to controls in adolescence (Milbocker et al., 2024). MRE was also sensitive to mechanical property changes following exercise in humans with multiple sclerosis and strongly correlated with memory performance scores, providing further evidence that MRE is a promising technique for intervention development (Sandroff et al., 2017). This study expands upon these findings by adding an environmental complexity component. While exercise alone enhances proliferation of neuronal precursors and increases white matter growth, environmental complexity following exercise extends the survival of newly generated cells and promotes cellular integration into established networks in rats (Gursky & Klintsova, 2017; Hamilton et al., 2012). Therefore, we expect rats that experienced this exercise and environmental complexity intervention to have long-term benefits to brain health which will be reflected in our mechanical property measures. We also quantify the density of cortical PNNs and compare them with MRE outcomes to help us better understand their role in mechanical property changes within the FASD brain. Ultimately, we present MRE as a potential method for noninvasively monitoring FASD progression and PNN integrity.

## 2. Methods

### 2.1 Study Design and Timeline

36 female Long-Evans rat pups (Charles River Laboratories) were divided equally into alcohol-exposed (AE) and sham-intubated (SI) treatment groups for longitudinal MRE scanning. We used a well-established rat model of FASD that mimics third trimester binge alcohol exposure in humans, coinciding with the brain growth spurt (Gursky & Klintsova, 2017; Milbocker & Klintsova, 2021). AE rats were administered 5.25 g/kg/day of ethanol in milk substitute via intragastric intubation twice daily during the rat brain growth spurt on postnatal days (PDs) four through nine (Dobbing & Sands, 1979). SI control pups received intragastric intubation without liquid administration to control for the stress of continual intubation during the brain growth spurt. Following intubations, pups were left undisturbed until weaning on PD 23 when they were separated into socially-housed (SH) groups of two to three rats per cage. Between PD 25-29, the first set of MRE scans was performed.

Following the first scan, half of the rats in each treatment group were randomly assigned to either the intervention combining wheel running and environmental complexity (WR/EC) or were socially-housed (SH) as controls. WR/EC rats were transferred to WR cages on PD 30 where they could freely exercise on an attached running wheel 24 hrs/day for 12 days. WR is an established method of modeling normal aerobic exercise (Patten et al., 2013; Van Praag et al., 2005). Control rats from each treatment group remained in SH cages from PD 30-42. Following WR, intervention-exposed rats were transferred to EC cages from PD 52-80. EC represents a combination of high social interaction, discovery, and learning (Diamond et al., 1964). It was modeled in this study using larger EC cages that housed six to nine rats and were furnished with various objects and toys changed daily for novelty (Gursky & Klintsova, 2017). Control rats remained in SH cages. Between PD 80 and 84, a second round of MRE scans was performed on all rats. Rodents were sacrificed immediately following their second MRE scans for brain tissue collection and PNN quantification. Final group sizes were: SI/SH = 9, SI/WR/EC = 9, AE/SH = 9, AE/WR/EC = 9. A separate cohort of rats was generated and sacrificed on PD 30 (n = 9 SI, n = 9 AE) to compare PNN measurements pre- and post-intervention, overlapping with the first and second MRE scanning time points. **Figure 1A** shows the timeline of this study.

**Figure 1.**
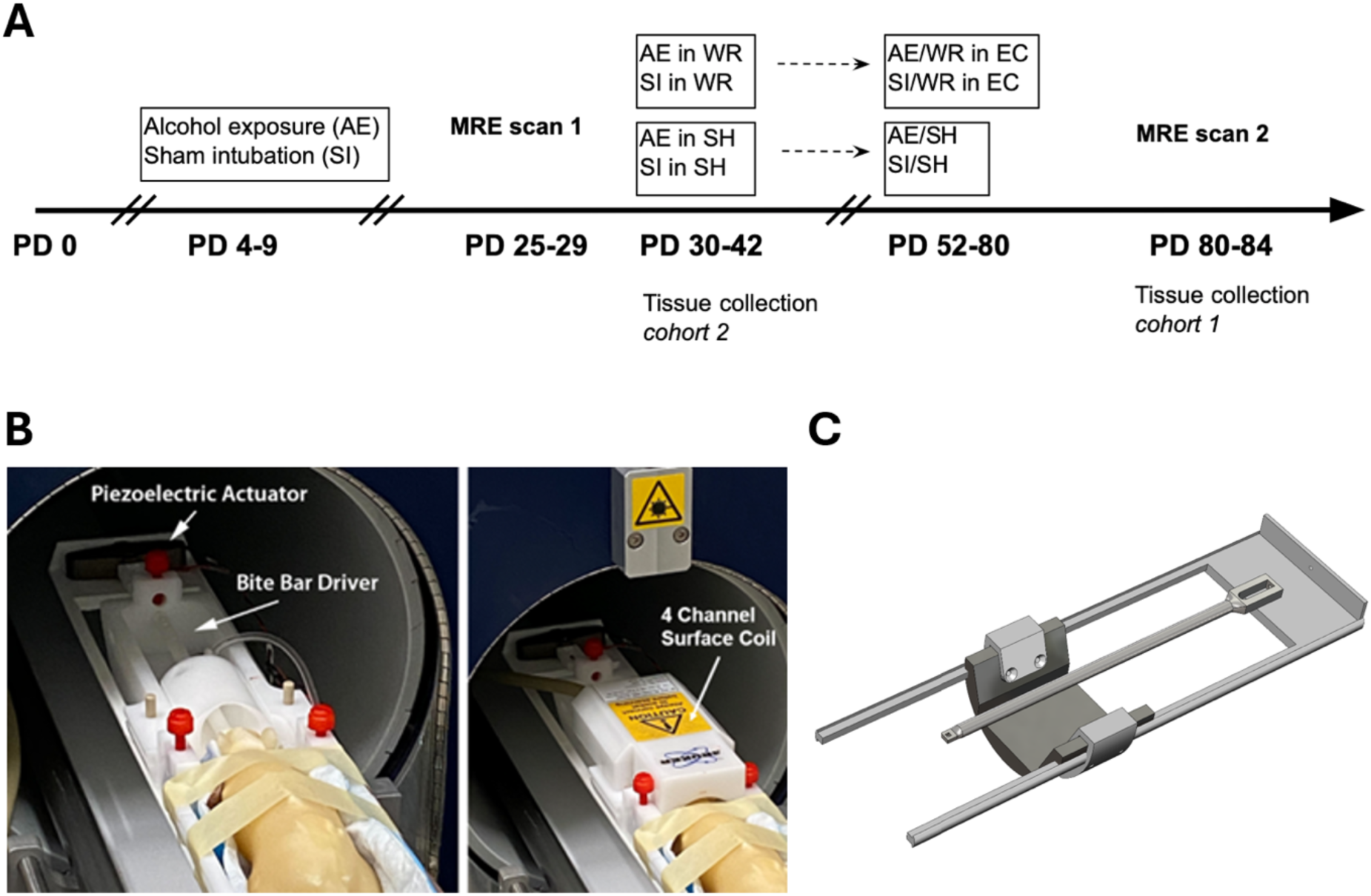
Study timeline and imaging setup. **(A)** Experiment timeline depicting establishment of the model of FASD (PD 4-9), WR/EC intervention (PD 30-80), pre- and post-intervention MRE scanning, and tissue collection. PD = postnatal day, AE = alcohol exposure, SI = sham intubated control, MRE = magnetic resonance elastography, WR = wheel running, SH = social housing, EC = environmental complexity. **(B)** Setup for MRE of the rat brain *in vivo* using a nonmagnetic, piezoelectric actuator, bite bar driver, and four channel surface receive coil. **(C)** 3D rendering of the actuator holder assembly. The actuator is mounted to the holder using a M2.5x0.45 screw. The brackets attach to the animal bed using two M3x0.5 screws on each side.

All procedures were performed in accordance with an approved University of Delaware Institutional Animal Care and Use Committee protocol and followed NIH ARRIVE guidelines for animal care. At no point in this experiment were rats single-housed.

### 2.2 Magnetic Resonance Elastography (MRE)

MRE scanning was performed on a Bruker 9.4T BioSpec preclinical MR imaging system (Bruker Corporation, Billerica, MA). Rats from all treatment and intervention groups were scanned pre-intervention, PD 25-29, and post-intervention, PD 80-84. Rats were anesthetized with 1-3% isoflurane administered via nose cone. Respiration and body temperature were monitored throughout the scan session, and temperature was regulated by the liquid-heated animal bed. 800 Hz vibrations were applied via a nonmagnetic, piezoelectric actuator (APA150M-NM, Cedrat Technologies, Meylan, France) attached to a custom 3D-printed bite bar resulting in average brain tissue displacement of 5-10 μm, following previously reported designs (Bayly & Garbow, 2018; Bigot et al., 2018; Milbocker et al., 2024). The actuator was mounted onto a custom 3D-printed bed extension which was secured to the animal bed with brackets on both sides. Guide rails designed to fit along tubing slots in the animal bed allowed for easy positional adjustments (**Figure 1B-C**). To maximize wave transmission, multiple bite bar sizes were designed and utilized to fit the rodents’ teeth snugly as they grew throughout development. Magnitude and phase images were acquired using the following imaging parameters: TE/TR = 60/3400 ms, FOV = 20 x 20 mm^2^, matrix = 80 x 80, slice thickness = 0.5 mm, slices = 40, averages = 24. A resolution of 0.25 x 0.25 x 0.5 mm^3^ was achieved in 32 minutes. Total scan time including other anatomical images was approximately one hour. Strong wave motion and data quality were confirmed via octahedral shear strain-based signal-to-noise ratio (OSS-SNR), and scans with OSS-SNR below three were excluded (McGarry et al., 2011).

Mechanical properties were estimated from displacement data collected during MRE scanning using the isotropic, nearly incompressible, nonlinear inversion (NLI) with a no boundary condition formulation (Kurtz et al., 2024; McGarry et al., 2012). NLI has been used extensively in human MRE and has been tuned for the rodent brain (Milbocker et al., 2024). It uses a heterogenous finite element model with subzone-based parallelization to minimize the difference between measurements and the computational model by iteratively updating maps of the complex shear modulus, 𝐺 ∗ = 𝐺’ + 𝑖𝐺’’, via the storage modulus, 𝐺’, and loss modulus, 𝐺’’. The storage and loss moduli were then used to calculate shear stiffness 𝜇 = 2|𝐺^∗^|^2^ / (𝐺’ + |𝐺^∗^|), and damping ratio, 𝜉 = 𝐺’’/2𝐺’ (Manduca et al., 2001; McGarry & Van Houten, 2008). Brains were masked from magnitude images using the SHERM automatic brain extraction tool (Y. Liu et al., 2021) and voxel-wise mechanical property maps of the global forebrain were generated. Slices at the extremities of the brain were excluded due to poor wave motion which results in inaccurate estimation of mechanical properties. Properties were reported as the median value in the region and outliers outside of 2 IQR were also excluded. The resulting sample sizes were SI/SH = 8, AE/SH = 6, SI/WR/EC = 8, and AE/WR/EC = 9.

### 2.3 Histological Examination of Perineuronal Nets

At the time of sacrifice, all rats were anesthetized with 4% isoflurane followed by an intraperitoneal injection of a ketamine/xylazine mixture. Rats were perfused transcardially with 0.1M heparinized phosphate buffered saline (PBS, pH = 7.20) followed by 4% paraformaldehyde in 0.1M PBS (pH = 7.20). Whole brains were extracted, post-fixed with a 30% sucrose in paraformaldehyde solution, and stored at 4℃ until sectioned in coronal plane at 40μm. Sections were stored in cryoprotectant solution maintaining anatomical order.

PNNs were visualized using either a nickel-enhanced diaminobenzidine (DAB) based chromophore (juvenile tissue collected on PD 30) or using a fluorophore (adult tissue collected on PD 80). In both cases, every eighth section in the rostro-caudal direction from anteroposterior coordinates -2.5mm to -5.0mm (Paxinos and Watson, 1982) was incubated with Wisteria floribunda agglutinin (WFA^+^, Vector Laboratories, B-1355-2, 1:100) overnight to identify PNNs. The following day, juvenile tissue was triple rinsed with 0.1M PBS before a 60-minute incubation in an avidin-biotin complex (ABC)-PBS solution (25 μL A and B to 12.5 mL of 0.1M PBS) followed by three rinses in 0.1M PBS. Tissue was submerged in a DAB solution in 0.1M PBS (0.05% DAB+0.01% hydrogen peroxide) for up to five minutes and then triple rinsed in 0.1M PBS. Adult tissue was triple rinsed with 0.1M TBS and then incubated in streptavidin AlexaFluor 647 conjugate (ThermoFisher Scientific, S-32357, 1:500) in TBS-NDS blocking solution for 120 minutes. When incubation was complete, the tissue was triple rinsed with 0.1M TBS for 10 minutes each.

Stained tissue was mounted, in anatomical order, on microscope slides and either coverslipped with gelvatol or, in the case of DAB-based staining, dehydrated and coverslipped with DPX mounting medium. The cortex was identified and outlined using a Paxinos and Watson rat brain atlas (7^th^ edition) and MBF Stereology software (MBF Biosciences). Fluorescently-stained tissue was imaged using a Zeiss AxioImager M2 microscope with Colibri 7 LED illumination (Carl Zeiss AG, Oberkocken, Germany) and brightfield images were acquired using a Zeiss Axioskop 2 Plus microscope.

Optical densitometry (OD) of PNNs in the cortex was measured using ImageJ. ImageJ calibration was performed using the Rodbard or Inverse OD functions. A step tablet containing a range of shades from white to black was uploaded, and the mean gray value of each shade was obtained. The corresponding OD value for each gray value was used to globally calibrate the program. TIF images containing the cortex were uploaded into ImageJ and transformed into 8-bit grayscale images. The cortex was outlined, and OD was measured for each brain section. The background OD of the section was also measured. The absolute value of the difference in OD (cortex-background) was obtained for each section and then averaged per animal to give one final OD score per subject.

### 2.4 Statistical Analyses

MRE measures were compared within-rat across time using repeated measures MANOVA with Greenhouse-Geisser correction for lack of sphericity. One-way ANOVAs were performed on MRE data collected on PD 25 to compare the effects of AE treatment with SI on stiffness and damping ratio. To comprehensively evaluate alterations to brain stiffness and damping ratio in adulthood between treatment and intervention groups, two-way MANOVA were performed. *Post hoc* analysis with Bonferroni correction for multiple comparisons was used to determine specific group differences. T-tests were also performed to test for average differences in mechanical properties between time points within corresponding postnatal treatment/intervention groups.

The absolute value of the difference in OD between the cortex and background was calculated to determine the average density of PNNs in the cortex (Gursky et al., 2020; Milbocker et al., 2023). OD for all sections containing the cortex were averaged within-animal to determine mean cortical OD per rat. Due to the nature of the fluorescent stain, the presence of perineuronal nets was represented by an increase in white space in the 8-bit grayscale TIF image. Therefore, the values are lower than those reported from brightfield staining where DAB deposits correspond to PNN presence.

A nonparametric independent samples test (Mann Whitney-U) was performed on PNN data collected on PD 30 to compare the effects of AE treatment with SI on PNN density. Two-way MANOVAs were performed to evaluate treatment x intervention exposure on PNN density on PD 80. *Post hoc* analysis with Bonferroni correction for multiple comparisons was used to determine specific group differences. Outliers outside of 2 IQR were discarded.

## 3. Results

Forebrain shear stiffness was significantly affected by alcohol exposure during the brain growth spurt. A repeated measures MANOVA evaluating treatment x intervention exposure on forebrain stiffness revealed significant effects of both time (*p* = 0.025) and treatment (*p* = 0.031), demonstrating that shear stiffness increased in both AE and SI rats over time from PD25 to PD 80 albeit shear stiffness increased more in AE rats compared to SI rats between time points. Post-hoc analyses revealed that juvenile AE rats exhibited softer forebrain tissue than SI controls on PD 25 (SI = 6.58 ± 0.48 kPa, AE = 6.17 ± 0.53 kPa, *p* = 0.031), but that the difference in stiffness between AE and SI rats on PD 80 was not significant, irrespective of intervention group (SI = 6.75 ± 0.53 kPa, AE = 6.58 ± 0.29 kPa, *p* = 0.307, **Figure 2**).

**Figure 2.**
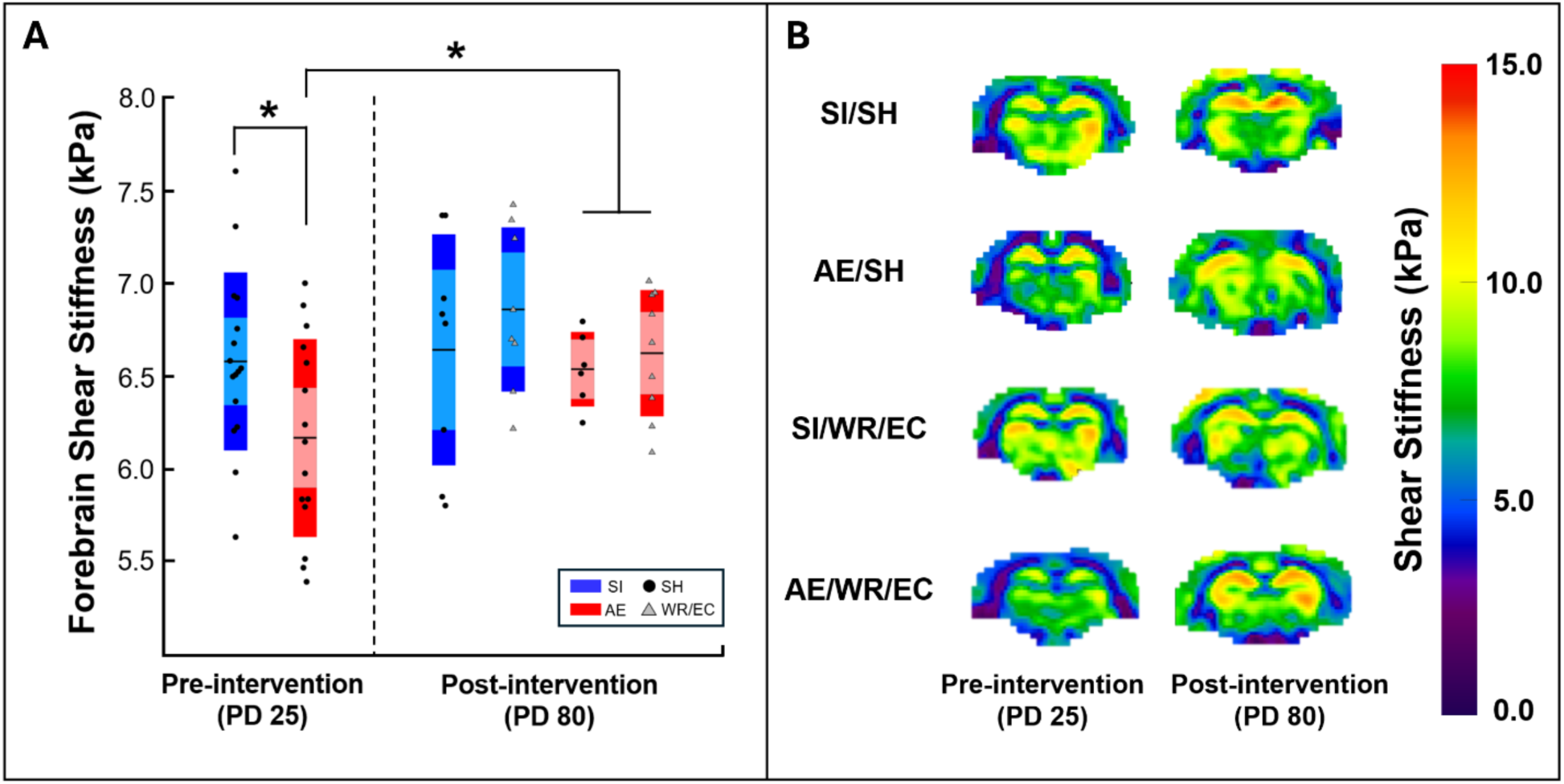
Forebrain shear stiffness in all rats in juvenility and adulthood. **(A)** Forebrain shear stiffness plot for each group across time. Alcohol-exposed rats (AE) had lower shear stiffness than controls (SI) pre-intervention and exhibited a larger increase in shear stiffness over time than controls. **(B)** Representative forebrain shear stiffness maps from each group collected on PD 25 and 80.

Damping ratio is a mechanical property that describes the relative viscous-to-elastic behavior of tissue. A repeated measures MANOVA evaluating treatment x intervention exposure on forebrain damping ratio revealed a significant time effect (*p* = 0.035) wherein damping ratio increased between PDs 25 and 80 in all rats, but there were no significant effects of treatment or intervention (**Figure 3**). Indeed, AE and SI rats exhibited similar damping ratios on PD 25 (SI = 0.209 ± 0.028, AE = 0.204 ± 0.028, *p* = 0.633) and PD 80 (PD 25: SI = 0.237 ± 0.048, AE = 0.225 ± 0.033, *p* = 0.401, **Table 1**).

**Figure 3.**
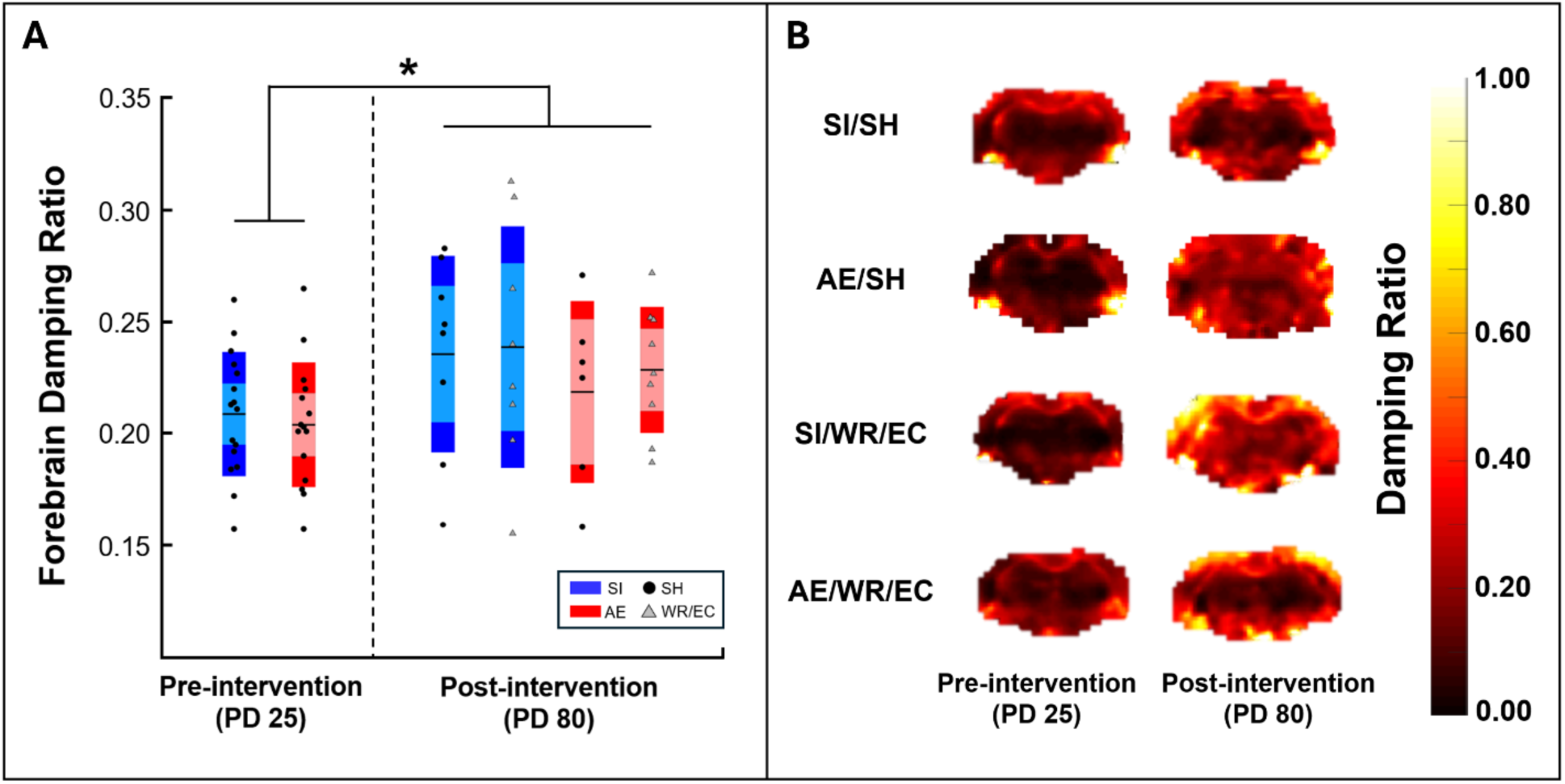
Forebrain damping ratio in all rats in juvenility and adulthood. **(A)** Forebrain damping ratio plot for each group across time. Both alcohol-exposed (AE) and sham-intubated (SI) control rats exhibited increases in damping ratio over time. **(B)** Representative damping ratio maps from each group across time.

**Table 1.** Summary of mechanical property results. Data are presented as mean ± standard deviation.

| PD 25 |  |  | PD 80 |  |  |
| --- | --- | --- | --- | --- | --- |
| Group | Shear Stiffness<br>(kPa) | Damping Ratio | Group | Shear Stiffness<br>(kPa) | Damping Ratio |
| SI | 6.58 ± 0.48 | 0.209 ± 0.028 | SI (combined) | 6.75 ± 0.53 | 0.237 ± 0.048 |
| AE | 6.17 ± 0.53 | 0.204 ± 0.028 | AE (combined) | 6.58 ± 0.29 | 0.225 ± 0.033 |
|  |  |  | SI/SH | 6.64 ± 0.62 | 0.236 ± 0.044 |
|  |  |  | SI/WR/EC | 6.86 ± 0.44 | 0.239 ± 0.054 |
|  |  |  | AE/SH | 6.54 ± 0.20 | 0.219 ± 0.041 |
|  |  |  | AE/WR/EC | 6.62 ± 0.34 | 0.229 ± 0.028 |

Nonparametric comparison of the OD of cortical PNNs on PD 30 revealed a significant effect of postnatal treatment on PNN density. In juvenile rats with AE, PNN density was significantly lower than in SI control rats (0.261 ± 0.096 vs. 0.375 ± 0.116; *p* = 0.028, **Figure 4A**), indicating that AE during the brain growth spurt significantly hinders PNN development in the cortex (**Figure 4B**). We did not observe any effect of treatment or intervention on PNN density on PD 80 (SI/SH: 0.133 ± 0.026, SI/WREC: 0.119 ± 0.017, AE/SH: 0.122 ± 0.015, AE/WREC: 0.104 ± 0.015; *p* = 0.500, **Figure 4C**), mirroring our MRE results.

**Figure 4.**
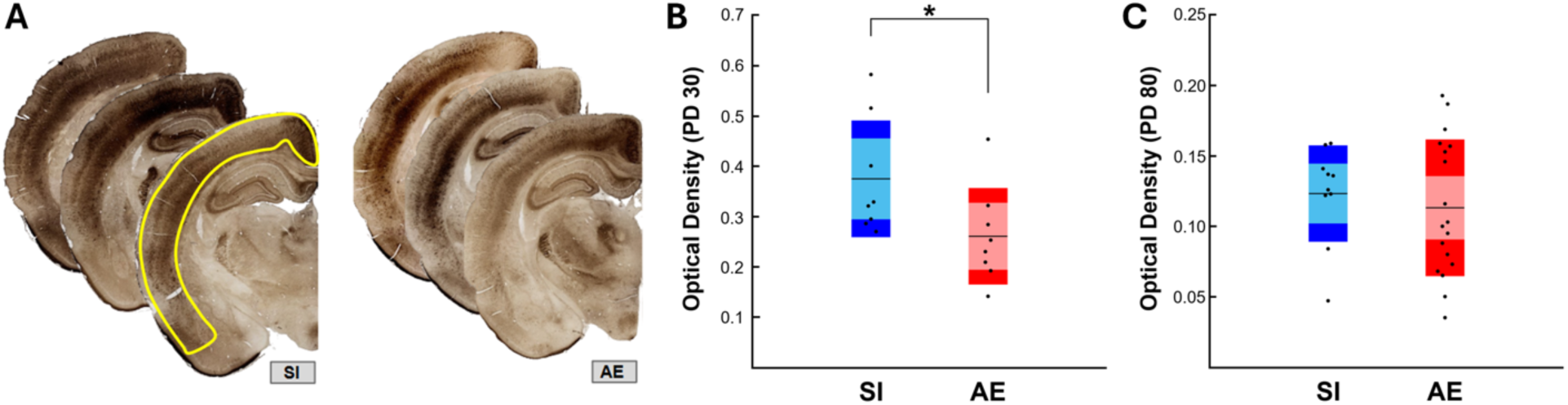
Histological results. **(A)** Micrographs of WFA^+^ brain tissue with cortex outlined in yellow from anterior to posterior (back to front). The density of WFA^+^ PNNs is greater in SI rats compared to AE rats in juvenility (PD 30). **(B)** Graph depicting the average optical density of WFA^+^ PNNs in the cortex at PD 30 (visualized with DAB). **(C)** Graph depicting the average optical dens WFA^+^ PNNs in the cortex at PD 80 (visualized with a fluorescent stain). Data from PD 80 were collapsed across intervention groups. The scales of the y-axes reflect the difference in staining techniques used to visualize PNNs. Outliers outside of 2 IQR were discarded (3 total), and error bars represent SD.

## 4. Discussion

In this study, we utilized MRE to demonstrate that the mechanics of the brain are altered during the juvenile period (PD 25-29) compared to young adulthood (PD 80-84) following prenatal alcohol exposure in a rat model of FASD. Mechanical properties increased over time and were affected by alcohol exposure during the brain growth spurt (PD 4-9). Shear stiffness was reduced in alcohol-exposed rats compared to controls in juvenility, and recovered to normal levels by adulthood. Conversely, damping ratio, a measure of tissue viscosity, was unaffected by postnatal treatment but did increase in all rats between PD 25 and 80. We anticipated that alcohol or age-related changes to the extracellular matrix might contribute to forebrain stiffness and damping ratio. We found that PNN density in the cortex, a region with the highest number of parvalbumin interneurons and documented sensitivity to prenatal alcohol exposure in humans and animal models, was significantly reduced in AE compared to SI control rats in juvenility. Moreover, PNN density did not differ between treatment and intervention groups in adulthood on PD 80. These findings appear consistent with the observed shear stiffness changes, suggesting that MRE could be a valuable tool for assessing nuanced changes to neurodevelopmental trajectory in juveniles with FASD. Ultimately, this may prove valuable to intervention development by establishing PNNs as a novel target for pharmaceutical therapy and by demonstrating how MRE may be a useful tool for assessing therapeutic efficacy in this population.

As expected, alcohol-exposed juvenile rats exhibited lower forebrain shear stiffness than sham intubated controls. This agrees with similar findings from animal models of other neuropathologies. Lower shear stiffness is often indicative of poor tissue health due to the impact of microstructural damage or delayed development on mechanical properties. Shear stiffness is lower in rodent models of Alzheimer’s disease (Munder et al., 2018; Murphy et al., 2012), multiple sclerosis (Millward et al., 2015; Wang et al., 2020), and stroke (Freimann et al., 2013). Our previous study also observed reduced forebrain stiffness in adolescent female AE rats compared to SI controls (PD 45, Milbocker et al., 2024). In both the previous and current studies, reductions in forebrain stiffness in AE rats were resolved by young adulthood (PD 75-85), independent of intervention exposure. These congruent results suggest that neurological damage due to prenatal alcohol exposure imposes a transient developmental delay in forebrain stiffening during which the juvenile-adolescent period is susceptible to stunted maturation. This is consistent with the existing neuroimaging literature on juveniles and adolescents diagnosed with Alcohol-Related Neurodevelopmental Disorder under the FASD umbrella that uses more conventional MRI scanning techniques (Hoyme et al., 2005; Inkelis et al., 2020; Milbocker et al., 2022, 2023; Riley et al., 1995; Sowell et al., 2008; Treit et al., 2013; Wozniak et al., 2019).

However, our knowledge of how development affects rat brain mechanical properties is limited as only a few MRE studies on development exist (Johnson & Telzer, 2018; Khair et al., 2023). One study found that juvenile rats with hydrocephalus exhibited differences in viscoelasticity compared to controls which was not observed in adult rats (Pong et al., 2017), indicating that brain and skull development mitigated the mechanical effects of hydrocephalus. In contrast, teratogenic exposure during gestational neurodevelopment (maternal immune activation with poly (I:C) on GD 15) prevented cortical stiffening in exposed offspring from juvenility into young adulthood (L. Liu et al., 2023). Importantly, Liu et al. did not observe developmental stiffening of deep gray matter structures between PD 28-70, irrespective of gestational treatment, in direct contrast to existing rodent MRE studies on brain development and our results. In humans, children with cerebral palsy exhibit lower whole brain shear stiffness than typically developing children (Chaze et al., 2019). When comparing healthy children and adults, shear stiffness differences vary by brain subregion but no significant differences were found globally (McIlvain et al., 2018). Furthermore, a follow up study by McIlvain et al. showed that brain stiffness gradually decreases over time from childhood to adulthood (McIlvain et al., 2022). This contrasts with our findings in rats, which all experienced increases in shear stiffness over time, but does agree with findings in developing mice (Guo et al., 2019). One possible explanation for this discrepancy is that shear stiffness does increase in humans, just during infancy and prior to the ages studied, and that the mechanisms driving stiffness increase occurs over a wider age range in rodents.

Damping ratio has been reported less frequently than shear stiffness, and as such, the effects of neurological disease on damping ratio are less clear. The damping ratio is higher in children with cerebral palsy (Chaze et al., 2019) and is negatively correlated with cognitive function (Hiscox, Johnson, McGarry, Schwarb, et al., 2020; Johnson et al., 2018; Schwarb et al., 2019). Based on our previous findings in adolescent FASD rats, we expected juvenile AE rats to have lower damping ratio than SI controls (Milbocker et al., 2024). However, we did not observe any differences between the groups on PD 25. While we did not find differences in damping ratio between SI and AE rats at either time point, damping ratio increased across all rats over time. This agrees with findings in humans where the damping ratio gradually increases from childhood to adulthood (McIlvain et al., 2022), but disagrees with findings in mice which did not exhibit global changes in phase angle, an analog of damping ratio (Guo et al., 2019), across development. Our previous study found that damping ratio increased over time from PD 42 to PD 70, but only in AE rats. Considering both studies together, the differences in damping ratio effects with AE and development may be attributed to specific critical periods within adolescent brain development that occur between the time points studied. The broad interval from PD 25 to PD 80 in the current study may mask shorter, more dynamic developmental windows. Similar critical periods have been observed in other studies where specific deficits in fear learning behaviors emerge transiently during circumscribed developmental windows within adolescence, notably between PD 24-33 (Heroux et al., 2019; Robinson-Drummer et al., 2018). Ultimately, these studies highlight the need for more continued longitudinal developmental MRE research with additional scanning time points to best capture periods of neurodevelopmental maturity after childhood.

We quantified cortical PNN density to assess how extracellular matrix integrity related to whole brain MRE outcomes in this model. PNNs are highly specialized and organized components of the ECM that form mesh-like webs around interneurons and changes in the structure or quantity of PNNs could lead to restricted plasticity of interneurons and decreased inhibitory control (Coleman et al., 2014). We found that PNN density was reduced in AE rats compared to controls in juvenility, suggesting that PNN maturation is delayed when AE overlaps with PNN formation (PD 4-18, Carulli & Verhaagen, 2021). Given the role of PNNs as a key permissive factor controlling critical periods in neurodevelopment, as well as in neuronal and synaptic plasticity that could be altered by alcohol insult, the reduced PNN expression could indicate altered plasticity, which is a scenario repeatedly demonstrated after adolescent and adult AE (Chen et al., 2015; Chen & Lasek, 2020; Dannenhoffer et al., 2022; Lasek, 2016). Notably, studies of adolescent AE in rats and binge drinking in adult mice reveal a significant link between ethanol exposure and diminished behavioral flexibility and suggest a potential novel mechanism involving changes in the number of inhibitory neurons and expression of PNNs. Delayed maturation of PNNs in the cortex would contribute to characteristic impairments in inhibitory control observed in FASD as it is guided by thalamocortical interactions (Ferguson & Gao, 2018).

Importantly, these PNN changes appear to mirror mechanical property changes. Shear stiffness is lower in juvenile AE rats, and this is consistent with a lower density of cortical PNNs in the same anteroposterior portion of the brain. Here, we provide mounting evidence that noninvasive MRE is likely sensitive to alterations in the organization of the brain’s extracellular matrix, even in the context of neurodevelopmental disorder. Silva et al. were the first to compare PNN integrity with MRE outcomes in adult rodents with neuropathology (R. V. Silva et al., 2024). The authors demonstrated that progression of neuroinflammation in a model of experimental autoimmune encephalomyelitis was correlated with decreased global brain stiffness and PNN density in the cortex. It must be noted that PNNs are not the only factor contributing to mechanical property estimates. Bergs et al. (2024) described a reductionist multi-network model that considers four major cellular networks contributing to MRE outcomes: neurons (gray matter), glia (neuroimmune system and white matter), extracellular matrix (PNNs), and vasculature. In our previous study on FASD, we examined the effect of AE on white matter integrity in relation to FASD. We observed inconsistent relationships between mature oligodendrocytes and shear stiffness or damping ratio results in adolescence and young adulthood (Milbocker et al., 2024). In contradiction, Schregel et al. (2012) demonstrated that in a rodent model of Multiple Sclerosis, aberrant deterioration of white matter via apoptosis of oligodendrocytes was positively correlated with a reduction in brain stiffness in adult mice. This suggests that oligodendrocytes may not contribute significantly to global MRE measures in this model. Future studies involving MRE of models of neurodevelopmental delay will require comprehensive analysis of several components of the brain parenchyma, as outlined by Bergs and colleagues, in addition to further investigation of the morphology, reactivity, and organization of mechanosensitive cells.

The exercise-environmental complexity intervention utilized in this study was chosen because it has been reported to be more powerful than exercise alone in inducing neuroplasticity (Hamilton et al., 2012, 2014; Rosenzweig, 1966). Aerobic exercise increases neurogenesis while novel learning introduced through a complex environment stimulates synaptogenesis of new neurons, thereby proliferating and integrating newly generated cells into neural circuitry (Hamilton et al., 2012). In an effort to capture these long-term changes, we scanned rats prior to and immediately following the intervention. We previously found that voluntary running in adolescence had a positive effect on forebrain shear stiffness in AE rats, allowing them to “catch up” to sham-intubated control levels (Milbocker et al., 2024). Due to the integration and proliferation of cells via environmental complexity, we expected to measure higher shear stiffness in control and AE/WR/EC rats. No differences were found in our MRE measures or PNN density between any of the subgroups post-intervention. It is also possible that the intervention influenced mechanical properties, but it was masked by a larger developmental effect. This was apparent by significant differences in rodent size and stage of neurodevelopment between time points manifesting as large time effects regardless of intervention group. An additional scanning time point immediately following wheel running and before environmental complexity caging may have helped separate the effects of intervention and development. Furthermore, several rats had to be excluded from the study, resulting in unequal and smaller subgroup sizes than anticipated. If a small intervention effect existed, it may have required a larger sample size to be measured.

## 5. Conclusion

In summary, this study built upon previous findings and confirmed that MRE is sensitive to mechanical property changes in FASD due to both alcohol exposure effects and neurodevelopmental processes. Our results indicate that prenatal alcohol exposure impairs development of brain tissue structure, as assessed through shear stiffness from MRE. PNN density followed similar trends to mechanical properties, suggesting that MRE is a powerful tool for noninvasively monitoring PNN density in FASD progression and intervention development. In future work, we will be exploring our central hypothesis that prenatal alcohol exposure leads to aberrant brain mechanical properties during development, mediated by disruptions in PNN formation and maturation, and can be an important clinical tool in assessing the FASD brain and monitoring response to therapies.

## Author Contributions

**LT Williams:** conceptualization, data curation, formal analysis, investigation, methodology, writing – original draft

**KA Milbocker:** conceptualization, data curation, formal analysis, investigation, methodology, writing – original draft

**E Smith:** conceptualization, investigation, writing – review and editing

**CP Bodner**: data curation, formal analysis, investigation, writing – review and editing

**DA Caban-Rivera:** data curation, investigation, writing – review and editing

**MDJ McGarry:** methodology, resources, writing – review and editing

**EEW Van Houten:** methodology, resources, writing – review and editing

**AY Klintsova:** conceptualization, project administration, resources, writing – review and editing, funding acquisition, supervision

**CL Johnson:** conceptualization, methodology, project administration, resources, writing – review and editing, funding acquisition, supervision

